# Derivation of human extended pluripotent stem cells in feeder-free condition

**DOI:** 10.1101/2020.10.18.344259

**Authors:** Ran Zheng, Ting Geng, Dan-Ya Wu, Tianzhe Zhang, Hai-Nan He, Liyan Wang, Haining Du, Donghui Zhang, Yi-Liang Miao, Wei Jiang

**Affiliations:** Department of Biological Repositories, Frontier Science Center for Immunology and Metabolism, Medical Research Institute, Zhongnan Hospital of Wuhan University, Wuhan University, Wuhan 430071, China; Institute of Stem Cell and Regenerative Biology, College of Animal Science and Veterinary Medicine, Huazhong Agricultural University, Wuhan 430070, China; State Key Laboratory of Biocatalysis and Enzyme Engineering, School of Life Science, Hubei University, Wuhan 430062, China; Hubei Key Laboratory of Cell Homeostasis, College of Life Sciences, Wuhan University, Wuhan 430071, China; Human Genetics Resource Preservation Center of Wuhan University, Wuhan 430071, China; Hubei Provincial Key Laboratory of Developmentally Originated Disease, Wuhan 430071, China

**Keywords:** extended pluripotency, totipotency, human pluripotent stem cell, epigenetic regulation, metabolic reprogramming

## Abstract

Human extended pluripotent stem cells (EPSCs), with bidirectional chimeric ability to contribute to both embryonic and extra-embryonic lineages, can be obtained and maintained by converting embryonic stem cells (ESCs) using chemicals. However, the transition system is based on inactivated mouse fibroblast, which greatly hinders the mechanistic studies of extended pluripotency and further applications. Here we reported a Matrigel-based feeder-free method to convert human ESCs and iPSCs into EPSCs and demonstrated the extended pluripotency in terms of molecular features, chimeric ability, and transcriptome. We further improved the protocol by applying chemicals targeting glycolysis and histone methyltransferase. Altogether, our data established a feeder-free system to generate human EPSCs and provided additional insights into the acquisition of extended pluripotency.

## INTRODUCTION

There are two distinct types of cells during mammalian early embryonic development: totipotent cells that harbor the superior development potential and are able to give rise to the whole conceptus including embryonic and extra-embryonic tissues, and pluripotent cells that can only contribute to embryonic lineages composed of most of the organs. The blastomere before the morula stage is considered as totipotent, which gradually loses totipotency and converts to pluripotency during the first lineage specification of embryo ^[1]^. Naive and primed pluripotency, two different metastable pluripotent states, have been detailly deciphered *in vivo* or their counterparts in vitro. Scientists have put numerous efforts into the conversion of primed human embryonic stem cells (ESCs) into a naive state ^[2–4]^, but not for totipotent state yet. Whether totipotent cells could be stably maintained *in vitro* or not was thus a long-standing question and poorly studied, particularly for human totipotent-like cells ^[5]^. Macfarlan and colleagues in 2012 first reported a transient 2-cell-like state existing in mouse ESC cultures, which exhibited similar transcriptome profile to *in vivo* 2-cell embryo and superior developmental potency to contribute to both embryonic and extraembryonic tissues ^[6]^. However, after the culture of these sorted 2-cell-like cells, they spontaneously turned into an ESC mix containing only less than 5% of 2-cell-like cells. In 2017, Yang and colleagues identified a cocktail medium called LCDM, which could culture mouse and human pluripotent stem cells into a different pluripotent state, named “extended pluripotency” ^[7]^. The extended pluripotent stem cells (EPSCs) exhibited outstanding developmental potential and bidirectional chimeric ability, contributing to both embryonic tissues and extra-embryonic tissues including yolk sac and placenta. Another group also independently established a chemical cocktail to maintain mouse single 8-cell stage blastomere *in vitro* as “expanded potential” stem cells ^[8]^ and later also succeeded in pig and human species ^[9]^. These two pioneering works give rise to irreplaceable cell types compared with naive or primed pluripotent stem cells in terms of both developmental potential and research significance. However, both of the conversion conditions were based on feeder cells, which brought uncertain factors to interfere with further molecular dissection and potential clinical application ^[10]^.

Here, we reported an optimized feeder-free culture condition to convert conventional human ESCs to EPSCs. We characterized the transcriptome profiles during the conversion and found the EPSCs exhibited more similarities to naive rather than primed ESCs, but still different from naive state. Human EPSCs expressed positive but relatively lower levels of core pluripotent genes including OCT4 and NANOG, and highly expressed genes enriched in zygotic genome activation, and more importantly, human EPSCs exhibited superior chimeric ability that contributed to both embryonic and extra-embryonic lineages. We further improved the protocol by applying chemicals targeting glycolysis and histone methyltransferase.

## RESULTS

### Generation of human EPSCs in feeder-free condition

Since the previous report that human EPSCs were established based on inactivated mouse embryonic fibroblast would hinder further mechanistic studies ^[10]^, we attempted to convert human ESCs into EPSCs without the feeder. Finally, we found LCDM plus another two chemicals IWR-1-endo and Y27632 (called LCDM-IY) with Matrigel were able to convert human ESCs into dome-shaped EPSCs. When human ESC line HUES8 was transferred to LCDM-IY medium, extensive cells started to differentiate and die, although very few maintained colony morphologies. Later, domed colonies that were distinct from flat conventional ESC colony emerged from passage 2 (Figure 1A). We hand-picked those dome-shaped colonies for several passages until domed colonies steadily appeared around passage 20. Then, the feeder-free EPSCs (called ffEPSCs) were maintained as domed colonies for more than 100 passages by single-cell dissociation every 2-3 days and followed by re-seeding at a low split ratio of 1:10 (Figure 1A).

**Figure 1.**
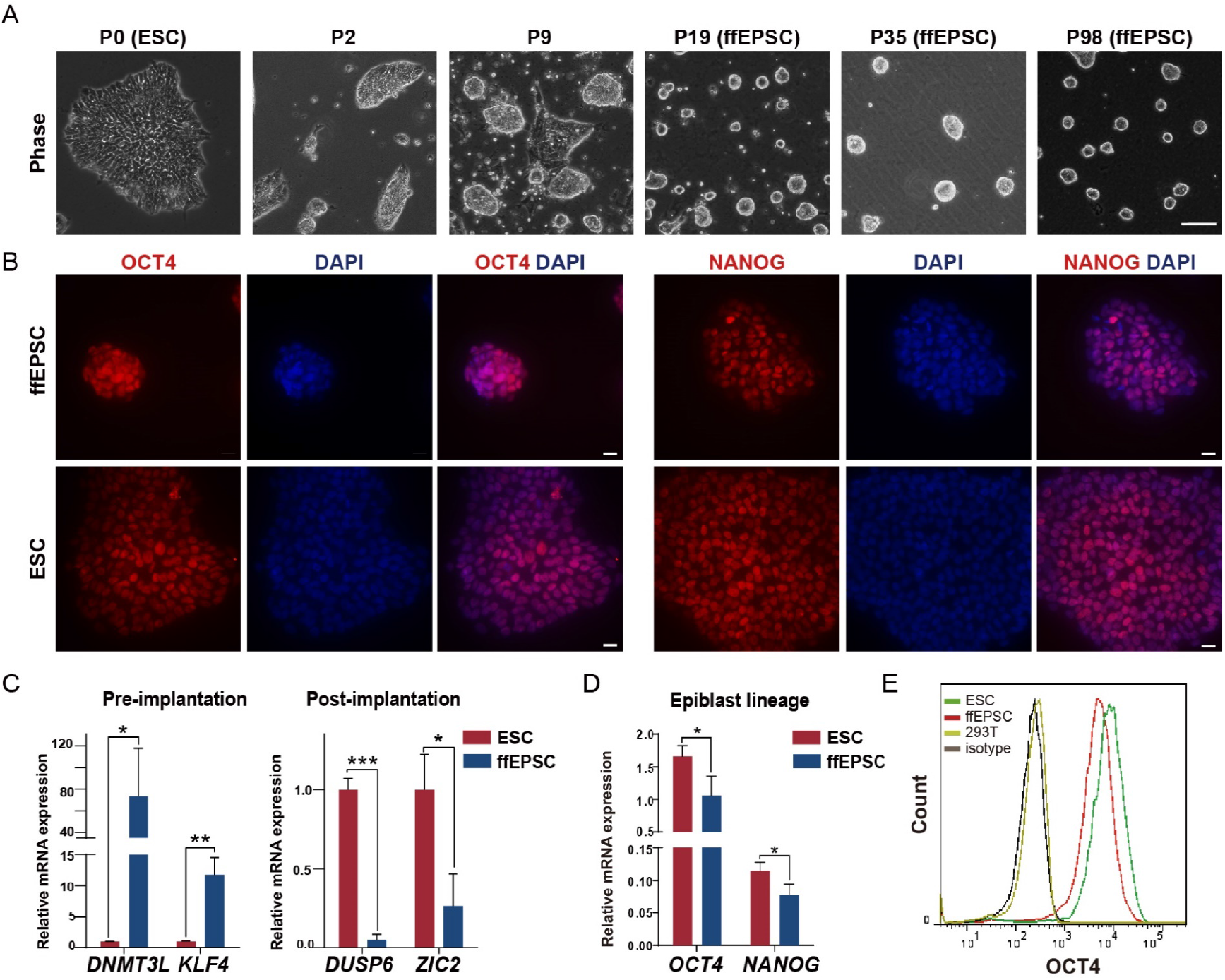
Generation of human EPSCs from ESCs under feeder-free condition. A. The morphology of cells during the transition of human ESCs into ffEPSCs. Scale bars = 100 μm. B. human ffEPSCs showed positive staining of NANOG and OCT4. ESCs served as positive control. Scale bars = 20 μm. C. Expression patterns of pre-implantation genes and post-implantation genes in ffEPSCs compared to ESCs (n=3, * p<0.05, ** p<0.001, *** p<0.001). D. RNA expression levels of *OCT4* and *NANOG* in ffEPSCs and ESCs analyzed by RT-qPCR (n=3, * p<0.05). E. human ffEPSCs showed lower signal of OCT4 compared to ESCs by flow cytometric analysis (n=3).

To characterize the pluripotency of converted ffEPSCs, we first checked the expression level of key pluripotent marker OCT4 and NANOG, and the ffEPSCs indeed maintained positive OCT4 and NANOG expression (Figure 1B). Since conventional human ESCs are considered as primed pluripotency rather than naive pluripotency, we further confirmed the expression levels of pre-implantation and post-implantation genes, as naive and primed markers respectively ^[2–4,11]^. By RT-qPCR analysis, we found ffEPSCs did express significantly higher pre-implantation genes such as *DNMT3L* and *KLF4*, but did not or rarely expressed post-implantation markers including *DUSP6* and *ZIC2* (Figure 1C). These data suggested ffEPSCs are much closer to earlier naive pluripotent state than later primed state. By RT-qPCR analyzing, we also found the epiblast lineage genes *OCT4* and *NANOG* were downregulated in ffEPSCs (Figure 1D). Since the EPSCs maintained the positive expression of core pluripotent genes *OCT4* and *NANOG* (Figure 1B), we further performed flow cytometry analysis to compare the protein levels. The result showed the protein level of OCT4 was decreased compared to that in conventional ESCs (Figure 1E), which was consistent with the notion that OCT4 was dispensable for the establishment of totipotency ^[12]^.

To determine the robustness of the feeder-free method, we applied our conversion system for multiple pluripotent stem cell lines. The results of another human ESC line H9 and one human iPSC line PGP1 ^[13]^ demonstrated the LCDM-IY feeder-free system was supportive to generate human ffEPSCs from various pluripotent stem cell lines. Similarly, many cells died or differentiated when transferred to LCDM-IY system, but a few cells survived and formed dome-shaped colonies (Figure S1A, D). These ffEPSCs could be stably maintained from around passage 16 and positively expressed OCT4 and NANOG (Figure S1B, F). Further RNA analysis suggested these ffEPSCs derived from different pluripotent stem cells highly expressed naïve markers rather than primed markers (Figure S1C, E).

### Transcription analysis of human ffEPSCs indicates similarity to human early embryo

To further investigate the molecular characters of ffEPSCs, we performed RNA-seq analysis for ffEPSCs and the parental ESCs. Compared with ESCs, the genes downregulated in ffEPSCs significantly enriched in developmental genes and cell differentiation-related genes, by looking into the different categories of developmental genes, we found that ffEPSCs expressed higher levels of early development genes and lower levels of embryonic developmental genes including ectodermal, endodermal and mesodermal genes (Figure 2A). And the up-regulation genes in ffEPSCs were noteworthily associated with epigenetics, DNA modification and histone modification (Figure 2A-B). Since the lower expression pattern of germ layer genes is a character of naive pluripotency compared to primed pluripotency, we further performed the gene set enrichment analysis (GSEA) and more interestingly, human zygotic genome activation genes are significantly upregulated in ffEPSCs (Figure 2C).

**Figure 2.**
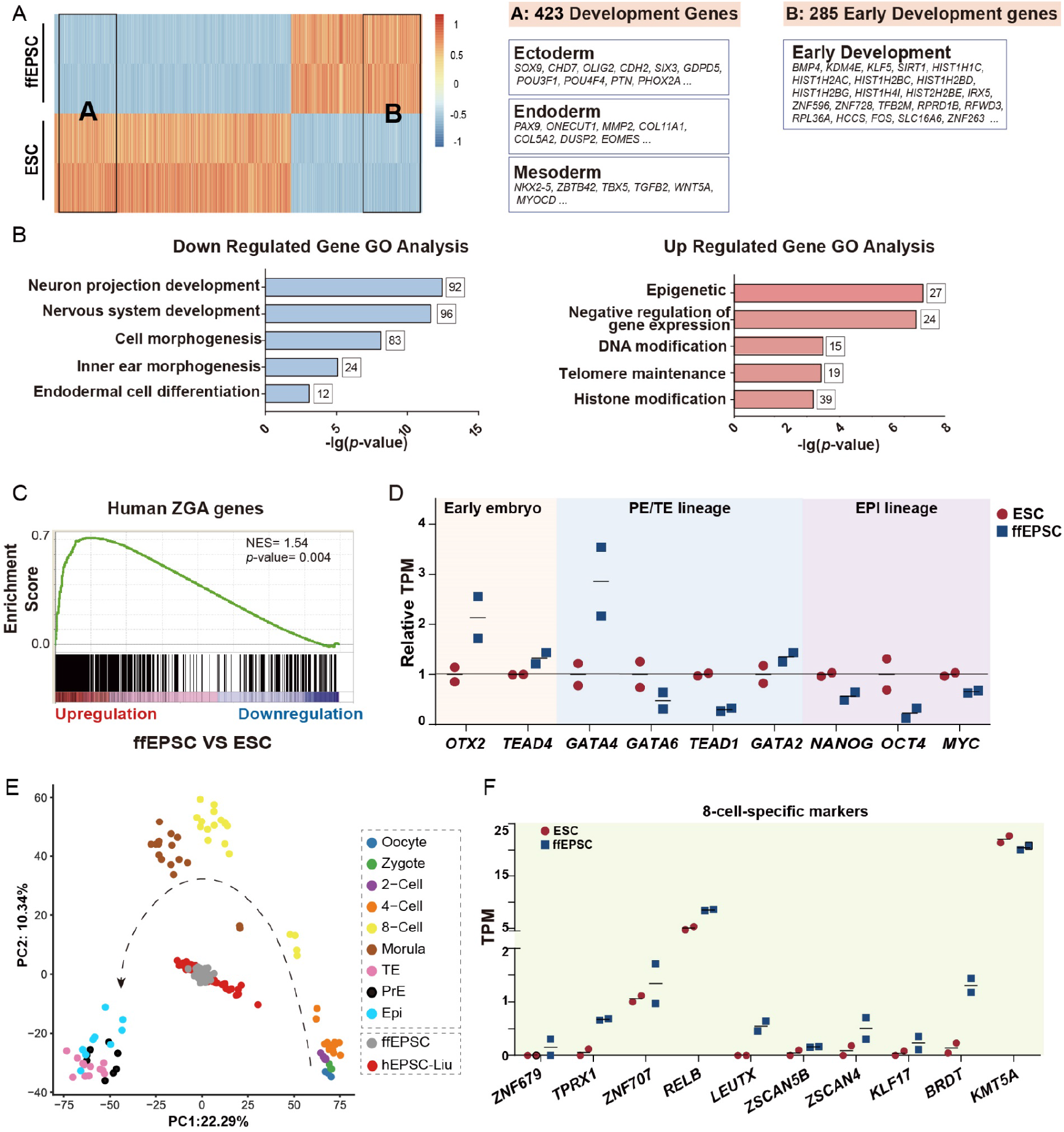
Transcriptome analysis of human ffEPSCs. A. Differential expressed transcripts between ffEPSCs and ESCs revealed by RNA-seq and developmental genes listed as different categories. B. Gene ontology analysis of the up- and down-regulated genes in ffEPSCs. C. GSEA analysis showed the expression pattern of human zygotic genome activation (ZGA) genes in human ffEPSCs and ESCs. D. Expression patterns of representative early embryo-specific genes, and primitive endoderm and trophectoderm (PE/TE) genes in ffEPSC and ESC. E. PCA and comparison of gene expression of single-cell RNA-seq dataset of human H1-EPSCs and human pre-implantation embryos showing ffEPSC more similar to 8-cell and morula stage. ffEPSCs, n = 25; oocyte, n = 3; zygote, n = 3; 2-cell blastomere, n = 6; 4-cell blastomere, n = 12; 8-cell blastomere, n = 20; morula, n = 16; late blastocyst, n = 30; hEPSC (from Liu group), n = 96; n represents the number of cells. F. Expression patterns of 8-cell-specific genes, identified by Stirparo and colleagues, in ffEPSC and ESC.

Furthermore, we found early embryo specific genes such as *OTX2* ^[14]^ and *TEAD4*^[15]^ were highly expressed in ffEPSCs, while primitive endoderm and trophectoderm lineage genes (*GATA4* and *GATA6*, *TEAD1* and *GATA2*, respectively) ^[16]^ were rarely expressed or downregulated (Figure 2D), suggesting an earlier stage of ffEPSCs than naive state. Consistent with this notion, we also found the epiblast lineage genes (*OCT4*, *NANOG* and *MYC*) were downregulated in ffEPSCs (Figure 2D) and RT-qPCR analysis confirmed the decreased expression levels of *OCT4* and *NANOG* in ffEPSCs (Figure 1D). We further performed single-cell RNA-seq of our ffEPSCs (n=25) as well, and then compared to the reference single-cell RNA-seq dataset about early embryo samples^[17]^, using the same protocol as Liu group for EPSC analysis^[18]^ (https://github.com/dbrg77/pig_and_human_EPSC). The result showed that global transcriptome of ffEPSCs located closer to 8-cell or morula stages, rather than late blastocyst samples including trophectoderm, primitive endoderm and epiblast lineages, similar to EPSC-Liu samples (Figure 2E). In addition, Stirparo et al. generated ten genes as the specific markers of 8-cell stage based on a number of RNA-seq datasets ^[19]^. We checked our data, and indeed nine of those ten 8-cell-specific genes showed higher expression in ffEPSCs compared with parental ESCs (Only *KMT5A* exhibited a little bit lower expression while highly expressed in both ESC and EPSC samples) (Figure 2F). Moreover, the higher expression levels of human zygotic activation genes and lower levels of *OCT4* and *NANOG* suggested ffEPSCs might represent an earlier pluripotent state different from primed or naive ESCs because early embryo of human and primate did not express OCT4 or NANOG until blastocyst stage ^[16]^. These data together support the conclusion that ffEPSCs exhibit closer transcription pattern to human early embryos.

### EPSCs exhibited the bi-directional chimeric ability

The fundamental characteristic of totipotent-like cell is the bidirectional chimeric ability. Therefore, we first constructed a human ESC line with stably expressed GFP reporter under the control of EF1a promoter, and then converted this reporter line to dome-shaped ffEPSCs (Figure 3A). We microinjected the single GFP-labeled ffEPSC into 8-cell-stage mouse embryos (Figure 3B) followed by 36-48 hours’ culture *in vitro*, and GFP signals were observed in embryos entering blastocyst stage (Figure 3C; 16 over 62 checked embryos), suggesting ffEPSC could support early embryo development. Furthermore, we performed immunostaining for the chimeric blastocysts to check the identity of GFP-positive cells. The result showed GFP co-stained with inner cell mass marker OCT4 or trophectoderm (TE) marker CDX2 or GATA3 ^[20]^ (Figure 3D) within the same blastocyst. Among the 19 GFP-positive blastocysts we checked, 9 showed co-staining of both GFP+/OCT+ and GFP+/TE+, 8 showed co-staining of GFP+/OCT+ only, and 2 showed co-staining of GFP+/TE+ only (Figure 3E). In addition, we also observed the co-staining of GFP with the extraembryonic endoderm marker GATA6^[21]^ in the chimeric blastocyst (Figure S2A).

**Figure 3.**
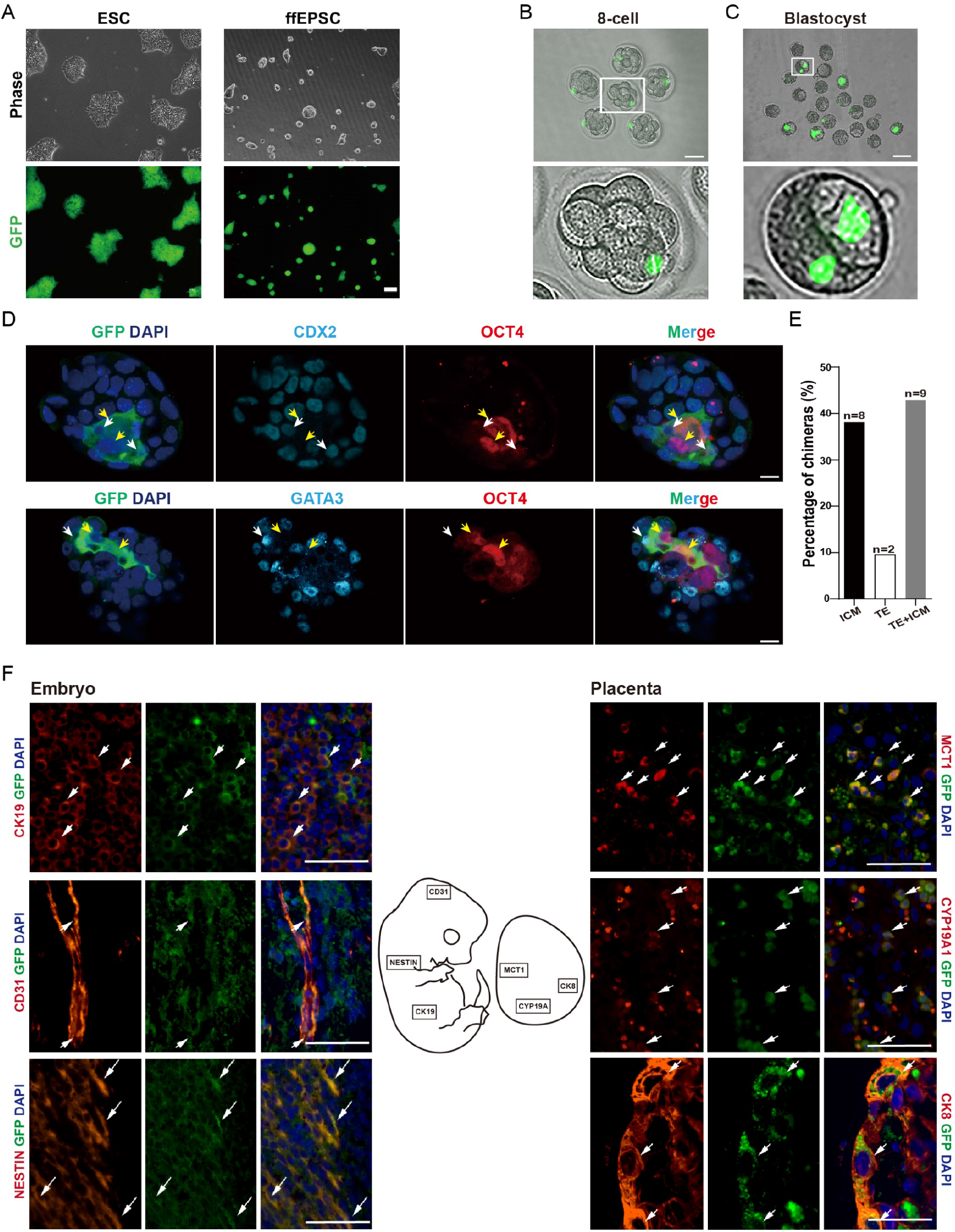
Human ffEPSCs exhibited bi-directional chimeric ability. A. Morphology and GFP fluorescence signal of ESC and ffEPSC. Scale bars = 100 μm. B. Mouse 8-cell embryos injected with single GFP-labeled ffEPSC. Scale bars = 50 μm. C. The mouse-human chimeric embryos developed to blastocyst stage, Scale bars =100 μm. D. GFP-labeled ffEPSC contributed to both inner cell mass (ICM, marked with OCT4) and trophectoderm lineage (TE, marked with CDX2 or GATA3) in mouse embryos. The arrows indicated the co-stained cells, Scale bars = 10 μm. E. Statistical results of bidirectional chimeric assay in D. F. Immunofluorescent staining of GFP (green) and lineage markers (red) in section of E13.5 embryo injected with GFP-labeled ffEPSC. NESTIN represents for ectoderm-derived neural tissue, CK19 represents for endoderm-derived gland tissue, CD31 represents for mesoderm-derived endothelial cells; and CK8 represents for pan-placental markers, MCT1 represents for syncytiotrophoblast I, and CYP19A1 represents for estrogen synthesis cells. Scale bars =25 μm.

We further determined the fate of GFP-positive cells in E13.5 chimeric embryos and placenta. The immune-fluorescence assay of GFP and lineage markers including NESTIN (ectoderm), CK19 (endoderm), CD31 (mesoderm) showed ffEPSC contributed to three germ layer lineages; meanwhile, GFP signal was found to co-localized with placenta markers CK8, MCT1^[22]^, and CYP19A1 (encoding aromatase for estrogen synthesis)^[23]^ (Figure 3F), indicating that ffEPSC could contribute to various placental cell lineages. We have also collected a series of embryos for DNA analysis using primer specific for human mitochondrial DNA and for mouse-human conserved DNA sequence, and the result shown below indicated ffEPSCs indeed contributed to both embryonic and extraembryonic (placenta and yolk sac) lineage in a certain extent at E13.5 stage (Figure S2B). Teratoma assay also supported this notion (Figure S2C). Taken together, the chimeric embryo analysis and teratoma assay showed that the converted ffEPSCs exhibited bidirectional chimeric ability and could differentiate into both embryonic and extraembryonic lineages *in vivo*, indicating the totipotency-like characteristic.

### Chemicals targeting glycolysis and histone methyltransferase facilitated the transition and maintenance of ffEPSCs

Based on the LCDM-IY-Matrigel system, we further explored the role of additional chemicals. Since a significant portion of the glycolysis-associated genes in ffEPSCs showed downregulation compared to ESCs (10/47, Figure S3A), but very few tricarboxylic acid cycle associated genes changed (Figure S3B), we treated ffEPSCs with glycolysis inhibitors including 2-deoxy-D-glucose (2-DG) and aurintricarboxylic acid (ATA)^[24]^. The results showed either 2-DG or ATA treatment groups increased the percentage of domed colonies although the number of colonies was similar (Figure 4A-B). The EPSCs treated with glycolysis inhibitors expressed higher expression levels of earlier stage genes such as *DNMT3L* and *KLF4* (Figure 4C). In addition, we performed the Seahorse flux analysis and the result showed the extracellular acidification rate (ECAR) level of EPSCs was significantly decreased in EPSCs (Figure 4D-E), indicating lower glycolytic activity in EPSCs compared to ESCs, which was consistent with the report that naive mouse ESCs displayed lower glycolysis level than primed EpiSC and human ESCs^[25]^. The dependence on oxidative metabolism supports a resemblance of these ffEPSCs to earlier embryo stages, and our data based on chemical inhibitors demonstrates distinct metabolism pattern may actively participate in cell fate determination. To further confirm the effect of glycolysis inhibitors, we chose iPSC-or H9-derived ffEPSCs for similar assay. 2-DG and ATA indeed increased the percentage of domed colonies yet with little effect on the number of colonies (Figure S3C-D, F-G). Supporting with this notion, 2-DG or ATA treated iPSC-derived ffEPSCs expressed higher *DNMT3L* and *KLF4* (Figure S3E, H). Those data together indicated that the application of glycolysis inhibitors improved the maintenance of ffEPSCs.

**Figure 4.**
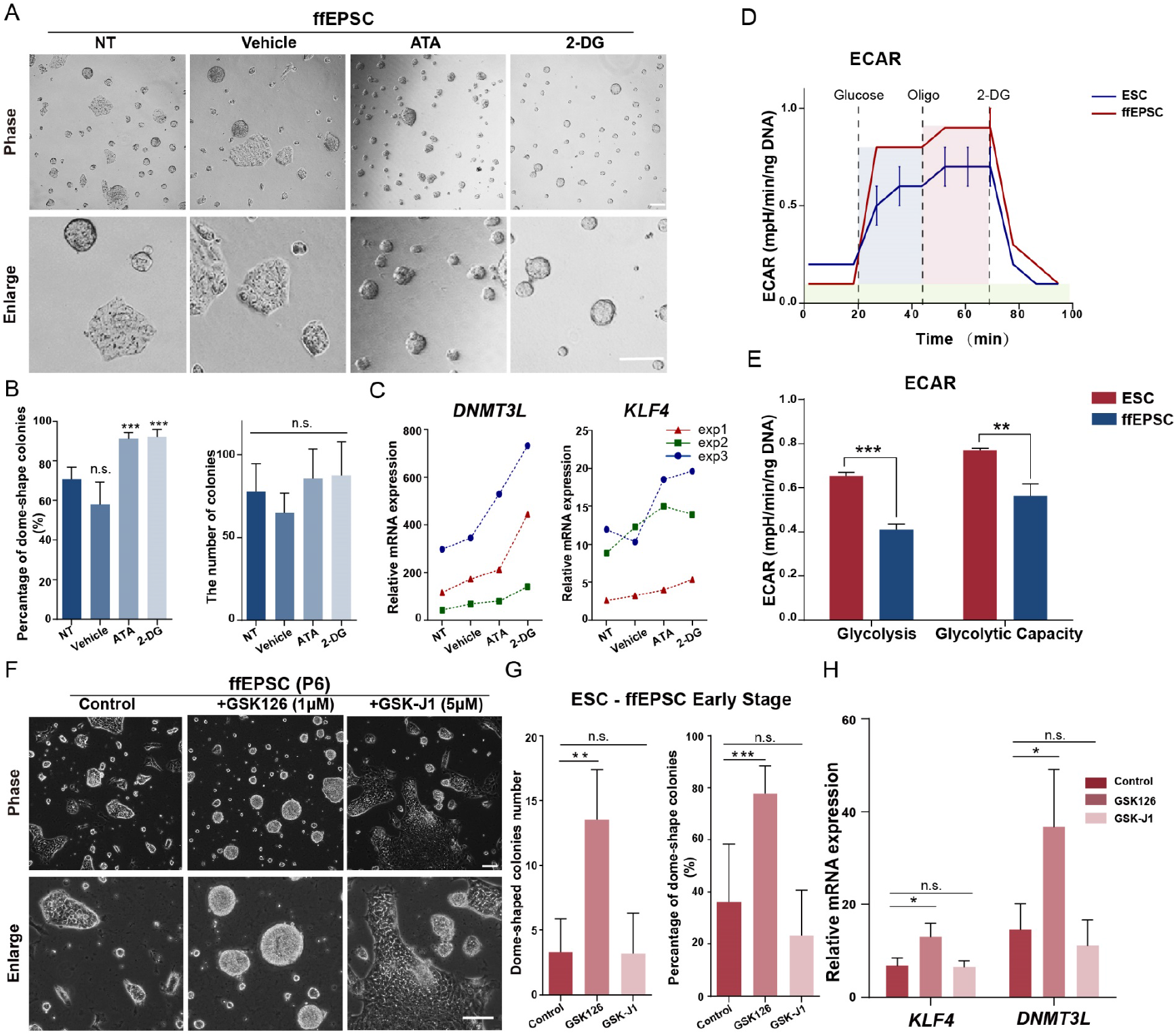
Chemicals facilitated the generation and maintenance of human EPSCs. **A.** Morphology of EPSCs cultured with different chemical inhibitors 2-DG and ATA. NT, non-treated. Scale bars = 100 μm. **B.** The number and percentage of dome-shaped colonies of EPSCs cultured with 2-DG or ATA(n=5, *** p<0.001). **C.** RT-qPCR analysis of pre-implantation genes after treatment with glycolytic inhibitors. Results from three independent experiments were shown. **D, E.** ECAR analysis of glycolytic rate and capacity of EPSCs measured by Seahorse flux compared to ESCs (n=3, ** p<0.01, *** p<0.001). **F.** Morphology of EPSCs cultured with different chemical inhibitors GSK-126 and GSK-J1. Scale bars = 100 μm. **G.** The number and percentage of dome-shaped colonies of EPSCs cultured with GSK-126, GSK-J1(n=3, ** p<0.01, *** p<0.001). H. Gene expression levels of EPSCs cultured with GSK-126, GSK-J1 or vehicle control (n=3, * p<0.05).

In addition, we investigated the effect of GSK126 targeting H3K27 methyltransferase EZH2^[26]^ and GSK-J1 targeting H3K27 demethylases^[27]^ during the EPSC transition. Interestingly, we observed the GSK126-treated group promoted more domed colonies and facilitated the transition (Figure 4F-G). Gene expression analysis also supported this notion that EZH2 inhibitor GSK126 facilitated EPSC-related genes expression (Figure 4H). Similar results were observed for H9-derived ffEPSCs (Figure S4A-B) and iPSC PGP1-derived ffEPSCs (Figure S4C). These data strongly supported that histone modification played an important role in acquiring human extended pluripotency from primed pluripotency. Further experiments would be absolutely needed to understand how H3K27 methylation regulated the transition.

## DISCUSSION

In this study, we started with optimizing the transition and maintenance condition of human EPSC in feeder-free, and then characterized the molecular and biological properties of the converted EPSCs in detail. Our data demonstrated the human ffEPSCs exhibited similar transcriptome to earlier embryo including relatively lower OCT4/NANOG level and upregulated zygotic activation gene set. Moreover, we further reported that glycolysis inhibitor and histone H3K27me3 methyltransferase EZH2 inhibitor could greatly facilitate the generation and maintenance of EPSCs, albeit the underlying mechanisms are waiting for further investigation.

Totipotent-like cell is a distinct cell category from pluripotent cell. First, EPSCs was recently shown to be able to greatly advance the efficiency in generating gene-targeting mouse models ^[28]^, indicating the significance and broad application. Second, OCT4 and NANOG are functionally expressed in both naive and primed pluripotent ESCs and iPSCs, but not or lower expressed in totipotent-like cells including either early human embryos before later morula stage ^[16]^ or 2-cell-like cells existing in mouse ESC cultures^[6]^. We observed ffEPSCs expressed OCT4 and NANOG, but maintained a relatively lower level compared to human ESCs (Figure 1D-E). However, it remains unknown how OCT4 and NANOG were downregulated in ffEPSCs, or even in early development. This pattern together with the transcriptome analysis (Figure 2) strongly indicated ffEPSCs exhibited a very distinct but an earlier state from naive or primed pluripotency. Herein, we believe that totipotent-like ffEPSCs can work as a key node between totipotency and pluripotency, and can serve as an advanced cell model to study early development at the molecular level in the future. The feeder-free condition to convert human pluripotent ESCs into EPSCs, established in this paper, should further promote the study of totipotency and application of EPSCs with superior chimeric ability.

Cellular metabolism is a complex and highly coordinated life-sustaining biochemical reaction that occurs within a cell that transforms or uses energy to maintain its survival. Several studies have analyzed the cellular metabolism patterns in naive and primed states. Naive mouse ESCs displayed lower glycolysis level than primed EpiSC and human ESCs ^[29]^. Consistent with this, here we revealed that human ffEPSC exhibited lower glycolytic activity than conventional human ESCs (Figure 4). Furthermore, distinct metabolism pattern may actively participate in cell fate determination. For instance, the ratio of α-ketoglutarate to succinic acid was critical to naive and primed pluripotency: high ratio of α-ketoglutarate to succinic acid promoted differentiation of human ESCs ^[30]^, but in mouse model, high ratio of α-ketoglutarate to succinic acid promoted mouse ESC self-renewal ^[31]^. In human ffEPSCs, here we reported the treatment of glycolytic inhibitors was beneficial to the maintenance of extended pluripotency. In addition, epigenetics serves as a pre-transcriptional factor that has been implied vital roles in development. Generally, global H3K27me3 serves as a repressive marker to alter the chromatin state in naive state pluripotency. Importantly, global H3K27me3 level shows dynamic remodeled in early stages blastomere, which is maintained in 2-to 8-cell stages, decreases at the 8-cell to morula stages and re-established in inner cell mass^[32]^. Moreover, histone modification was reported to play a critical role in the transition of mouse primed to naive state ^[33]^. Our data that chemical inhibition of methyltransferase facilitated the conversion of ffEPSCs and genetic depletion of demethylase blocked the conversion of ffEPSCs suggested H3K27me3 level served as a negative regulator in acquiring human extended pluripotency. In summary, our data provided a metabolic and epigenetic insight into the acquisition of extended pluripotency, which can help further understand the underlying mechanism.

## STAR★METHODS

### Maintenance and conversion of human ESCs to EPSCs

Human ESC line HUES8 was normally maintained on Matrigel (Corning)–coated (1:100) plates with mTeSR1 (STEMCELL Technologies) supplemented with 1% penicillin-streptomycin ^[34]^. For the conversion of ESCs to EPSCs, single cells treated by Accutase were replaced to Matrigel-coated 6-well plates with mTeSR1. On the next day, the medium was exchanged to the LCDM-IY medium. LCDM-IY medium was based on the mix of knockout DMEM/F12 and neurobasal medium (1:1), supplemented with 0.5×B27 Supplement, 0.5×N2 Supplement, 5% knockout serum replacement, 1% GlutaMAX, 1% nonessential amino acids, 1% penicillin-streptomycin, 0.1 mM β-mercaptoethanol, and additional six inhibitors including recombinant human LIF (10 ng/mL), CHIR99021 (1 μM), (S)-(+)-Dimethindene maleate (2 μM), Minocycline hydrochloride (2 μM), IWR-endo-1 (1 μM) and Y-27632 (2 μM). One or two days later, cells were collected by trypsin and re-seeded on Matrigel-coated plates (1:30) in LCDM-IY medium and cultured with daily change of the medium. Established EPSCs were passaged by TrypLE around every 3 days while conventional HUES8 were passaged by accutase every 5 days.

### Immunofluorescence staining

Cells were grown on 24-well plates and washed by DPBS before fixed with 4% paraformaldehyde for 20 minutes at room temperature. Fixed cells were blocked with PBS containing 10% donkey serum and 0.3% Triton X-100 for 2 hours at room temperature. Then cells were incubated in the blocking buffer with diluted primary antibodies at 4℃ overnight or at room temperature for 2 hours. The cells were further incubated in the blocking buffer with diluted second antibodies at room temperature for 2 hours after three times’ washing with DPBS. Nuclei were counterstained with DAPI (1:10000, Thermo-Fisher).

Mouse embryos were collected to 96-well plates with a round bottom, fixed with 4% paraformaldehyde for overnight at 4°C, and then permeated in PBS containing 0.5% Triton X-100 for 30 minutes and blocked with 10% donkey serum in PBS containing 0.3% Triton X-100 for 2 hours at room temperature. Three times’ washing with DPBS containing 0.01% Triton X-100 and 0.1% tween-20 was applied before every step. Embryos were incubated in the blocking buffer with diluted primary antibodies at 4℃ overnight, and then further incubated with diluted second antibodies at room temperature for 1 hour after washing. At last, embryos were put in mounting medium with DAPI for image capture under the Zeiss LSM880 microscope.

The antibodies used were listed as follows: Anti-NANOG mouse IgG (SantaCruz, SC-293121, 1:200), Anti-OCT4 mouse IgG (SantaCruz, SC-5279, 1:200), Anti-OCT4 rabbit IgG (Cell Signaling Technologies, 2750, 1:200), Anti-GATA3 rabbit IgG (Cell Signaling Technologies, 5852, 1:200), Anti-GATA6 rabbit IgG (Cell Signaling Technologies, 5851, 1:200), Anti-OCT3/4 rabbit IgG (BD, 611203, 1:200,). Anti-aromatase rabbit IgG (ABclonal, A12684, 1:100), Anti-MCT1 rabbit IgG (ABclonal, A3013, 1:100), Anti-CDX2 rabbit IgG (ZSGB-bio, ZA0520, 1:200), Donkey-Anti-Mouse-TRITC (Jackson immuno Research, 715-025-150, 1:200), Donkey-Anti-Mouse-488 (Jackson immuno Research, 715-545-150, 1:200), Donkey-Anti-Rabbit-TRITC (Jackson immuno Research, 711-025-152, 1:200), Donkey-Anti-Rabbit-FITC (Jackson immuno Research, 711-095-152, 1:200).

### Flow cytometry

Human ESCs were dissociated with accutase and EPSCs were dissociated with TrypLE, to generate single-cell suspension. Intracellular flow cytometry was operated with the Transcription Factor Buffer Set (BD Biosciences). Cells were stained with primary and then secondary antibodies diluted with 2% FBS in PBS containing 0.3% Triton X-100. Data were collected on a FACS Celesta flow cytometer (Becton Dickinson) and analyzed using FlowJo.

### RT-qPCR

Total RNA was extracted with the HiPure Total RNA Mini Kit (Magen). 1-2 μg of total RNA was reverse-transcribed into complementary DNA with 5×qRT super Mix. qPCR was done in duplicated with 2×SYBR Green qPCR Master Mix (Bio-Rad). Glyceraldehyde-3-phosphate dehydrogenase (GAPDH) was used as an endogenous housekeeping control. Student’s t-test (two-tailed, equal variance) was performed to obtain p-values for RT-qPCR experiments. Primer sequences are listed as follows: *NANOG* (CCCCAGCCTTTACTCTTCCTA, CCAGGTTGAATTGTTCCAGGTC); *OCT4* (CAAAGCAGAAACCCTCGTGC, TCTCACTCGGTTCTCGATACTG); *KLF4* (ACCCACACAGGTGAGAAACC, ATGCTCGGTCGCATTTTTGG); *DNMT3L* (CGCCCCATGTAAGGACAAGT, ATCGGGTGCAATCAGGGTTT); *ZIC2* (GCACGTCCACACCTCCGATAA, TGGACCTTCATGTGCTTCCGCAG); *DUSP6* (TGGAACGAGAATACGGGCG, CTTACTGAAGCCACCTTCCA); *GAPDH* (AATGAAGGGGTCATTGATGG, AAGGTGAAGGTCGGAGTCAA). Human-specific mitochondrial element (CGGGAGCTCTCCATGCATTT, GACAGATACTGCGACATAGGGT); Human-mouse conserved mitochondrial element (GCTAAGACCCAAACTGGGATT, GGTTTGCTGAAGATGGCGGTA).

### Next generation sequencing and data analysis

Total RNA of human ESCs and EPSCs were prepared in duplicate with the HiPure Total RNA Mini Kit (Magen) according to the manufacturer’s instruction. Samples were sequenced on Illumina HiSeq X Ten PE150 at Annoroad Gene Technology Co. Ltd.

Sequencing reads were aligned to the human genome build hg38/GRCh38 with the HISAT2 ^[35]^. Raw counts were performed with FeatureCounts ^[36]^ using GENCODE v29 human gene annotation ^[37]^. Raw counts were normalized for total read counts using the size factors computed by the Bioconductor package DESeq2 ^[38]^. Differential expression analysis was performed using the default settings of DESeq2 with p-value of <0.05 and filtering out genes with TPM less than 1. To generate the heatmap for differential gene expression, TPM values were scaled relative to the mean expression of each gene across all samples in R (http://www.r-project.org/). The Gene Expression Omnibus (GEO) accession number for the RNA-seq raw data reported in this work is GSE137208 and expression values were shown as TPM in Table S1. Functional annotation of significantly different transcripts and enrichment analysis were performed with Clusterprofiler ^[39]^.

To quantify the specific genes for zygotic genome activation stage, two RNA-seq datasets of human early embryo development (GSE44183 ^[40]^ and GSE36552 ^[41]^) were aligned to the human genome with the HISAT2 aligner, and raw counts were normalized to TPM and filtering out genes with TPM less than 20. The ZGA-specificity score of each transcript was defined as follows: Score = meanA - (meanOther+2*sdOther). Where meanA is the mean expression of the samples in certain stage, and meanOther and sdOther are the mean and SD of the expression levels in the other samples, respectively. Therefore, a positive score indicated that the gene was expressed in a certain stage at a considerably higher level than in the rest of the stages. A gene with a score of >0.5 was considered as specifically expressed in a certain stage. The Single-cell RNA-seq Data was performed by SMART-seq2 protocol. The PCA analysis source code were generated with https://github.com/Teichlab/NaiveDE/tree/master/NaiveDE.

For Gene Set Enrichment Analysis (GSEA) we used normalized counts by DEseq2 as input. The ZGA specific genes were generated by the overlap of two RNA-seq datasets (GSE44183 and GSE36552).

### Seahorse cellular flux assays

XF24 Cell Culture Microplates were pre-coated with Matrigel 2 hours before cell seeding. Human ESCs and EPSCs were seeded onto Matrigel-coated plate and cultured for 6 hours. Then the culture media was changed by base media (unbuffered DMEM supplemented with 2 mM Glutamine, pH 7.3-7.4) 500 μl per well about 1 hour before the assay. Selective chemical inhibitors with proper concentrations were added during the measurements. Cells glycolysis stress test were measured using an XF24 Extracellular Flux Analyzer. All the data were normalized to DNA concentration calibrated by CyQuant™ Cell Proliferation Assay kit (Thermofisher).

## ACKNOWLEDGEMENTS

We would like to thank Dr. Lei Zhang for help in writing and discussion, Jing Lv, Ran Liu, Chengli Dou and other laboratory members for technical help and discussion. WJ was supported by grants from the National Key Research and Development Program of China (No. 2016YFA0503100), the National Natural Science Foundation of China (No. 91740102, 31970608), the Medical Science Advancement Program of Wuhan University (No. TFZZ2018053, TFJC2018005) and the Fundamental Research Funds for the Central Universities (2042018kf0256). DZ was funded by the Ministry of Science and Technology of China (National Science and Technology Major Project, Grant No. 2018YFA0109100) and the National Natural Science Foundation of China (No. 31871496).

## Authors’ contributions

Jiang and D. Zhang conceived the project, and designed the experiment together with Zheng and Geng. Geng and Zheng performed most of the bench experiments and T. Zhang analyzed the NGS data. Wu performed the chimera assay with the help from Wang and He, and Zheng and Geng performed the embryo staining and image capture. Du provided experimental material regarding histone modification analysis. Jiang and Miao supervised the project. Jiang, Geng, Zheng, D. Zhang and Miao wrote the manuscript. All authors contributed to and approved the final manuscript.

## DECLARATION OF INTERESTS

The authors declared no competing interests.

## SUPPLEMENTAL FIGURES LEGENDS

**Figure S1.**
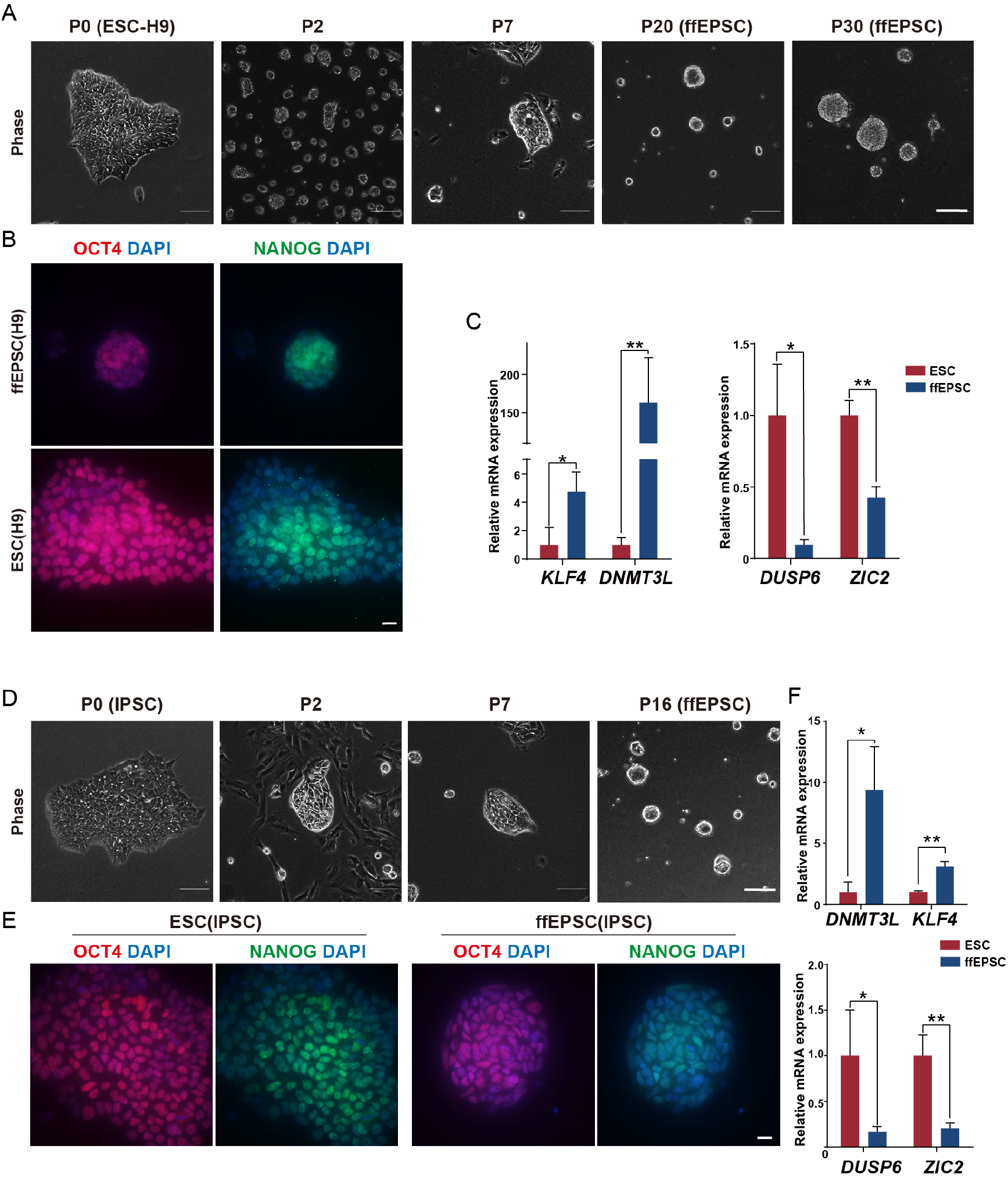
Generation of human EPSCs from H9 or iPSCs under feeder-free condition. A. The morphology of cells during the transition of ESC H9 into ffEPSCs. Scale bars = 100 μm. B. H9-derived ffEPSCs showed positive staining of NANOG and OCT4. Scale bars = 20 μm. C. Expression patterns of pre-implantation genes and post-implantation genes in ffEPSCs compared to H9-ESCs, data shown as average ±sd, n=3, * p<0.05, **p<0.01. D. The morphology of cells during the transition of human iPSCs into ffEPSCs. Scale bars = 100 μm. E. human iPSC-derived ffEPSCs showed positive staining of NANOG and OCT4. Scale bars = 20 μm. F. Expression patterns of pre-implantation genes and post-implantation genes in ffEPSCs compared to iPSCs.

**Figure S2.**
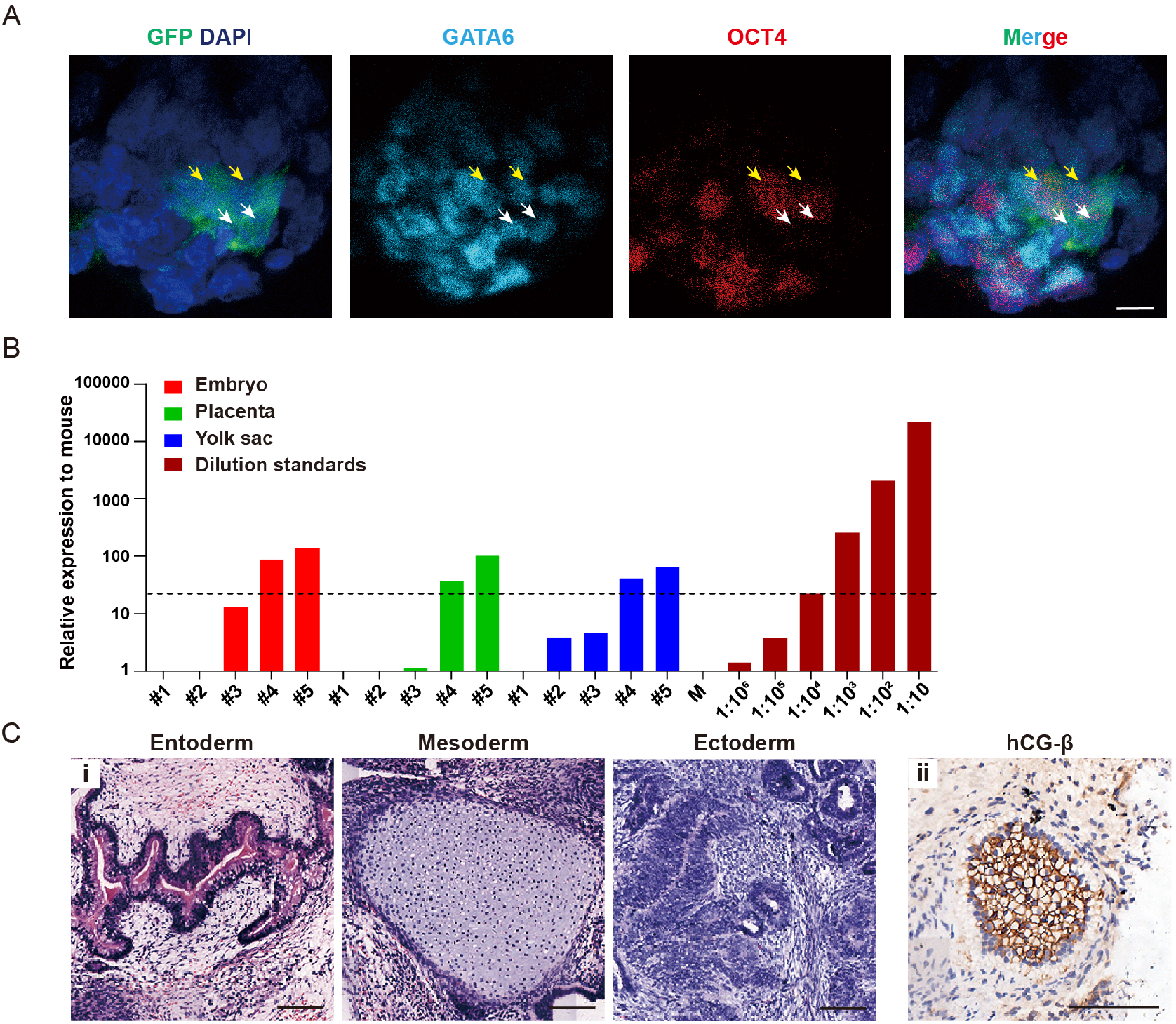
Human EPSCs harbored bi-directional developmental potential. A. EPSC-GFP contributed to both inner cell mass (ICM, marked with OCT4) and extraembryonic endoderm lineage (marked with GATA6) in chimeric mouse-human embryos. The arrows indicated the co-stained cells. Scale bars = 10 μm. B. Quantitative PCR measurements or human mitochondrial DNA indicated the presence of human cells in mouse embryos, placentas and yolk sac at E13.5 following injection of single ffEPSC at the 8-cell stages; a series of human-mouse cell dilutions were run in parallel to estimate the degree of human cell contribution. Black dotted line highlights level equivalent to 1:100,000 diluted standard. C. (i) Hematoxylin-eosin staining (for three germ layers) in teratoma section and (ii) Immunofluorescent staining of hCG-β (for extraembryonic lineage). Scale bars =100 μm.

**Figure S3.**
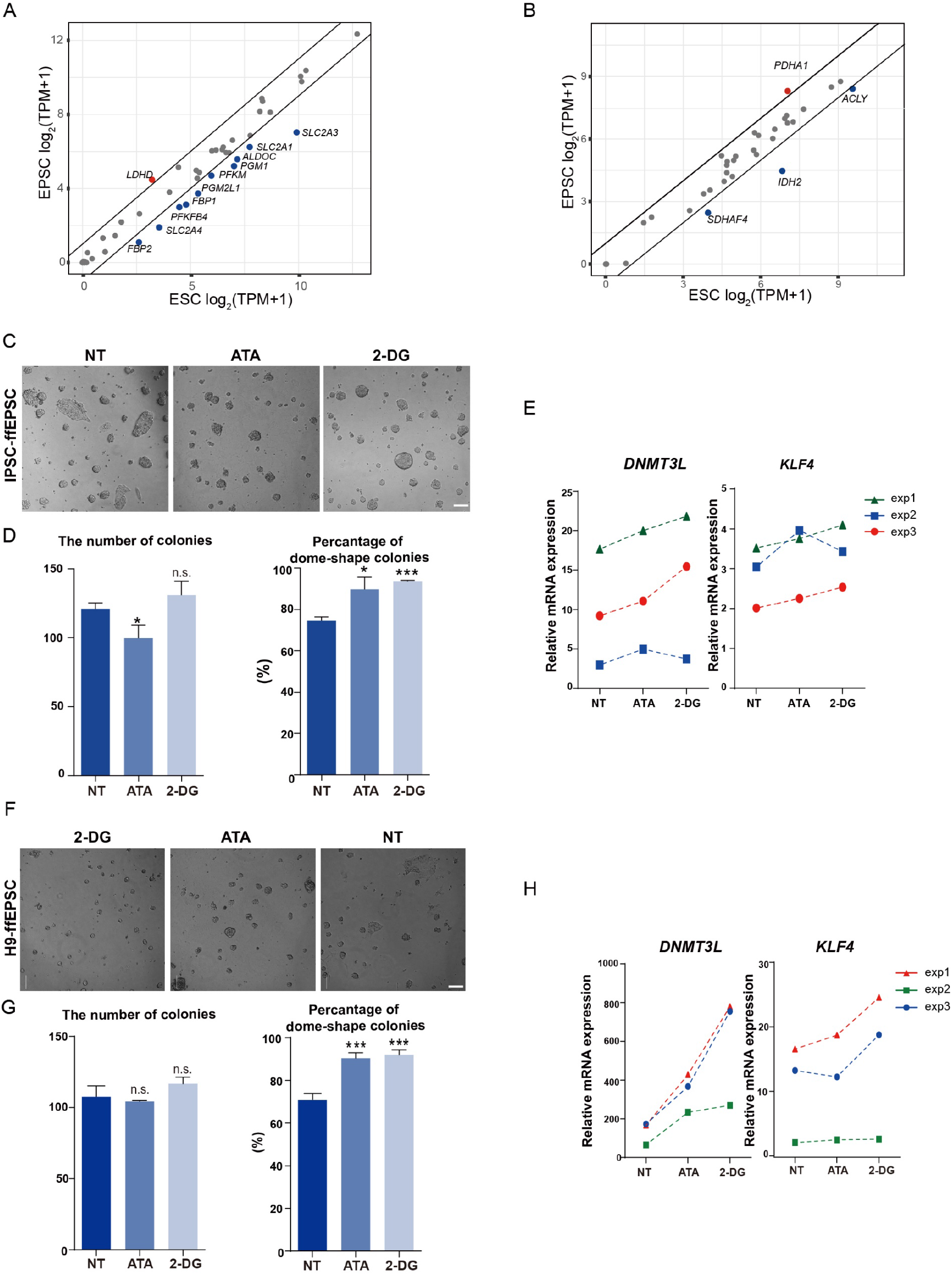
Glycolysis inhibitor 2-DG and ATA facilitated H9 or iPSC-derived human ffEPSC maintenance. A. The plot of expression levels of glycolysis-related genes in ESCs and ffEPSCs. The up-regulated genes are colored in red (1/47) and blue for downregulated gene (10/47). B. The plot of expression levels of TCA-related genes in ESCs and ffEPSCs. The up-regulated genes are colored in red (1/34) and blue for downregulated gene (3/34). C. Morphology of H9-derived ffEPSCs cultured with different chemical inhibitors 2-DG and ATA. Scale bars = 100 μm. D. quantitative number and percentage of dome-shaped colonies of H9-derived ffEPSCs cultured with different chemical inhibitors 2-DG and ATA, data shown as average ±sd, n=3, *p<0.05, ***p<0.001). E, RT-qPCR analysis of pre-implantation genes after treatment with glycolytic inhibitors. Results from three independent experiments were shown. F. Morphology of iPSC-derived ffEPSCs cultured with different chemical inhibitors 2-DG and ATA. Scale bars = 100 μm. G. quantitative number and percentage of dome-shaped colonies of iPSC-derived ffEPSCs cultured with different chemical inhibitors 2-DG and ATA, data shown as average ±sd, n=3, *p<0.05, ***p<0.001). H, RT-qPCR analysis of pre-implantation genes after treatment with glycolytic inhibitors. Results from three independent experiments were shown.

**Figure S4.**
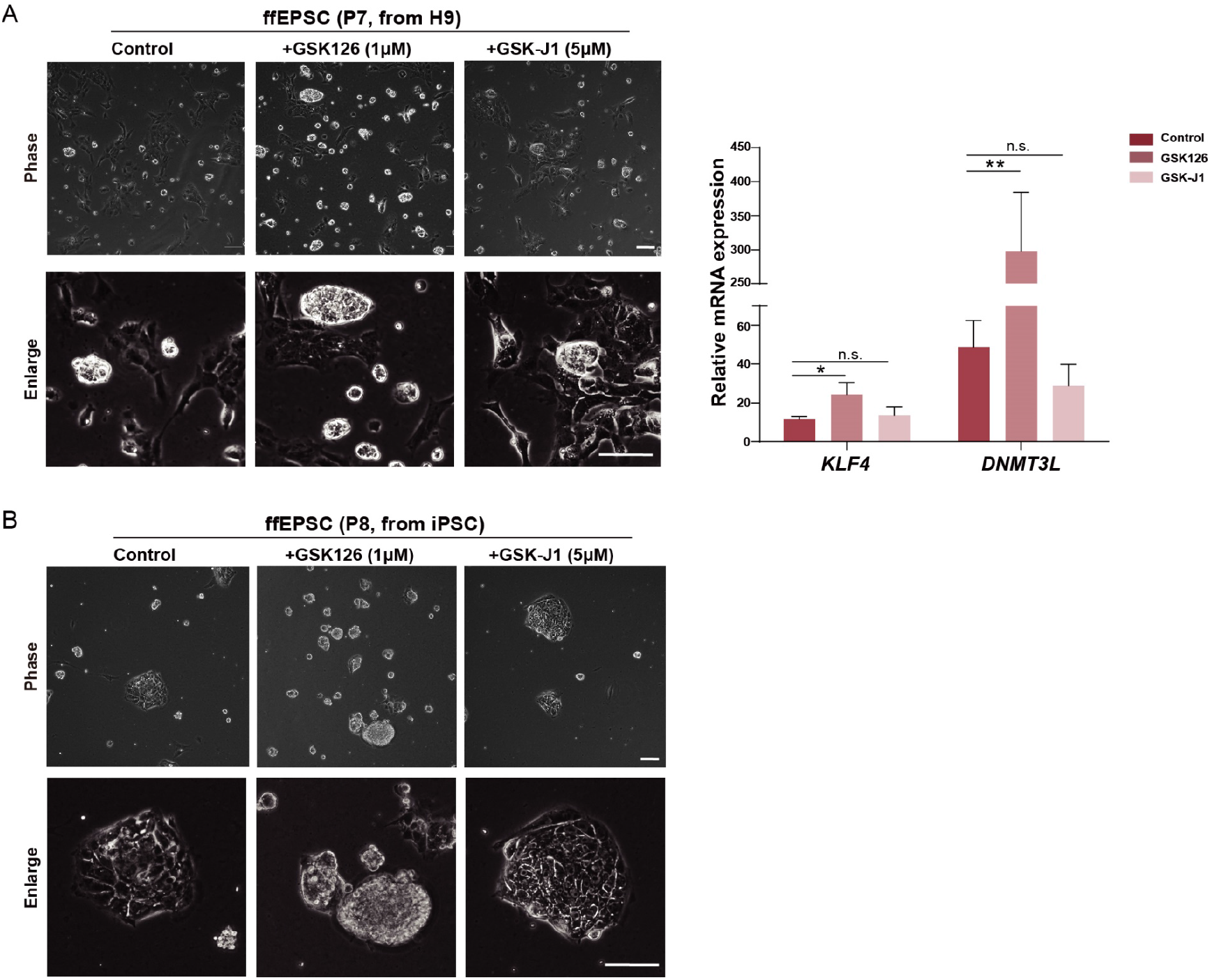
GSK126 facilitated human ffEPSC transition from ESC and iPSC. A, Morphology of H9-derived ffEPSCs cultured with different chemical inhibitors GSK126 and GSK-J1. Scale bars = 100 μm. B, Gene expression levels of ffEPSCs cultured with GSK-126, GSK-J1 or vehicle control (n=3, * p<0.05, **p<0.01) C. Morphology of iPSC-derived ffEPSCs cultured with different chemical inhibitors GSK126 and GSK-J1. Scale bars = 100 μm.

## Notes

### Competing Interest Statement

The authors have declared no competing interest.

